# DenMark: A Bayesian Hierarchical Model for Identifying Cell-Density Correlated Genes from Spatial Transcriptomics

**DOI:** 10.64898/2026.04.02.713482

**Authors:** Mingchi Xu, Alexandra Schmidt, Qihuang Zhang

## Abstract

Recent advances in single-cell-resolution spatial transcriptomics enable the profiling of gene expression while preserving the precise locations of individual cells, enabling quantitative investigation of how cellular organization relates to molecular state. A fundamental yet under-modeled aspect of organization is local cell density, which varies across microenvironments and can be linked to transcriptional programs. However, rigorous computational frameworks to quantify density-expression correlations remain lacking. Here, we present DenMark (<u>Den</u>sity-dependent <u>Mark</u>ed point process framework), a unified statistical framework that jointly models local cell locations and gene expression in single-cell-resolution ST data, enabling identification of density-correlated genes while naturally providing uncertainty quantification. To scale inference, DenMark leverages a Hilbert space Gaussian process approximation. In simulations, DenMark provides an accurate and well-calibrated estimate of density-expression association. Across single-cell ST platforms, including MERFISH and 10x Xenium, and across brain and cancer tissues, DenMark identifies genes whose expression is associated with cellular clustering and reveals density-related biological programs.

## Introduction

In multicellular systems, tissue and organ function emerge from the spatial organization of cells and the relative abundance of distinct cell populations (Rao et al., 2021). A key quantitative descriptor of this organization is cell density, the local abundance of cells per unit area or volume, which reflects the degree to which cells are packed within a microenvironment (O’Brien and Bilder, 2013). By influencing cell-cell communication (Youk and Lim, 2014), diffusion of signaling molecules (Su et al., 2024), and competition for shared resources (Levayer and Moreno, 2013), cell density provides a mechanistic link between tissue architecture and cellular state.

Mounting evidence suggests that changes in cell density are accompanied by coordinated transcriptional responses. In controlled *in vitro* settings, manipulating plating density or confluency alters the expression of gene programs across multiple cell types and culture systems (Wang and Adamo, 2000; Frisa and Jacobberger, 2002; Schmitt-Ney and Habener, 2004; McBride and Knothe Tate, 2008; Heng et al., 2011). Within *in vivo* tissues, density likewise varies systematically across microenvironments, for example, across tumor niches with immune infiltration (Oliveira et al., 2025), across developmental and anatomical layers (Maynard et al., 2021), or across regions of tissue remodeling (Agramunt et al., 2026), and these differences often coincide with distinct molecular states. Together, these observations motivate the study of *density-expression correlation*: how transcriptional variation relates to local changes in cellular abundance within local tissue contexts.

Recent advances in spatial transcriptomics (ST) provide an unprecedented opportunity to quantify density-expression correlation *in vivo*, which measures molecular features while preserving their spatial context within tissues. ST platforms include both sequencing-based approaches that profile transcriptomes from barcoded spots, and imaging-based approaches that directly detect RNA molecules *in situ*. Recently developed technologies, including seqFISH (Lubeck et al., 2014), MERFISH (Zhang et al., 2023), 10X Xenium (Janesick et al., 2023), and 10X Visium HD (10x, 2024), achieve the transcriptional profiling in single-cell or subcellular resolution, providing essential ingredients for investigating how local cellular density is associated with transcriptional variation.

Despite this opportunity, most statistical tools for ST primarily target the intrinsic spatial structure of gene expression rather than explicitly quantifying density-expression correlation. Existing approaches typically test for departures from spatial homogeneity or score spatial structure using coordinate- or graph-based neighborhoods, including using point-pattern and marked-point statistics (Edsgärd et al., 2018; Sun et al., 2020; Zhu et al., 2021; Bernstein et al., 2022), spatial Gaussian process frameworks (Svensson et al., 2018; BinTayyash et al., 2021; Weber et al., 2023), and graph-based or discrete spatial models (Hu et al., 2021; Zhang et al., 2022). These methods primarily aim to detect *spatially variable genes* (SVG), i.e., genes with spatially heterogeneous expression, rather than *density-correlated genes* (DCG), whose expression levels are statistically associated with local cellular density. These models often treat cell locations as fixed and model gene expression conditional on spatial coordinates or neighborhood structure, leaving variation in cell density unmodeled and providing limited uncertainty-quantified inference for the density-expression relationship.

To address this gap, we propose DenMark (<u>Den</u>sity-dependent <u>Mark</u>ed point process), a unified statistical inference framework that jointly models cell locations and gene expression in single-cell-resolution ST data to infer density-expression correlation with uncertainty quantification and facilitate the identification of DCGs. DenMark models where cell crowding occurs in space and how strongly each gene is expressed across that space, explicitly capturing shared spatial variation between local cell density and gene expression. The model decomposes spatial variation into components shared between density and expression and components that are gene-specific, providing an interpretable separation between density-driven transcriptional effects and residual spatial structure. To scale inference to high-resolution spatial transcriptomics data, DenMark employs a Hilbert space Gaussian process approximation (HSGP) (Solin and Särkkä, 2020). We validate DenMark using simulations, demonstrating accurate and computationally efficient estimation of density-expression correlation. In benchmarking analyses of MERFISH mouse brain datasets, DenMark recovers known density-correlated genes and identifies signals that are not captured by conventional SVG-focused approaches. Applying DenMark to a 10x Xenium breast cancer dataset, we identify density-associated transcriptional programs linked to spatial variation in immune infiltration, highlighting an antagonistic interplay between tumor and immune microenvironments.

## Results

### Workflow of DenMark

The general idea of DenMark is to quantify the correlation between the local cell density and the average gene expression per cell, and to identify density-correlated genes (DCGs) through a Bayesian hierarchical marked point process model (Figure 1). Here, DCGs represent genes whose expression levels vary with the local cell density (Figure 1A). For each candidate gene to be investigated, DenMark leverages the location and gene expression of each cell in spatial transcriptomics data to jointly model cell density and local average gene expression as two smoothly varying spatial processes. To approximate these spatially continuous processes, tissue samples are discretized into rectangular grids by aggregating the cell counts and gene expressions, respectively, in each grid (Figure 1B). These gridded counts and gene expression are then jointly modeled through a marked point process, where the spatial pattern of gene expression is decomposed as a common spatial component shared with the cell density and a specific spatial pattern solely presented in gene expression, revealing how gene expression per cell covaries with local cell density (Figure 1C). Parameters in DenMark are estimated following the Bayesian paradigm, and the resultant posterior distribution is approximated via Markov Chain Monte Carlo (MCMC) methods (Neal et al., 2011; Stan Development Team, 2024). As DenMark involves two Gaussian processes (GPs) and the computational burden scales with the size of the approximating grid, we use a Hilbert space Gaussian process (HSGP) to approximate them. This substantially reduces the computational cost of posterior inference. Finally, DenMark provides (1) a direct estimate of the correlation between cell density and local per-cell gene expression with uncertainty quantification, (2) a summary of the latent field decomposition to visualize how the information is shared between cell density and gene expression, and (3) a visualization of how spatial correlation between two locations decays as their distance increases (Figure 1D).

**Figure 1.**
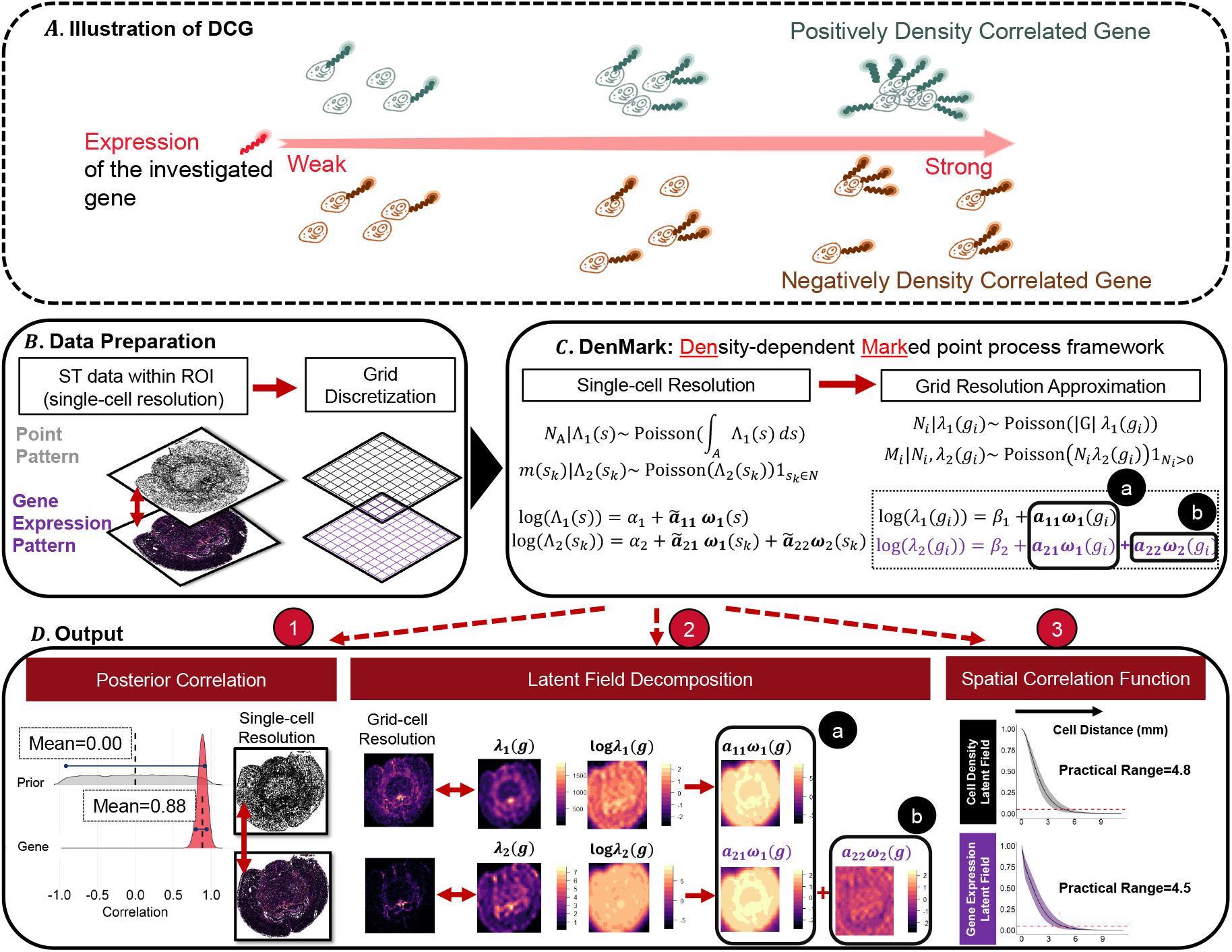
Workflow of DenMark. (A) Conceptual visualization of density-correlated genes. In regions of higher cell density, positively density-correlated genes show increased expression (upper panel), whereas negatively density-correlated genes show decreased expression (lower panel). (B) Data preparation: For each candidate gene, DenMark extracts cell location (point pattern) and cell-level gene expression measurements (gene expression pattern) from single-cell resolution ST data. These two spatial patterns are then discretized onto a set of predefined grids. (C) DenMark framework: The discretized grid-level cell counts and expression are fitted by the marked point process, facilitated by a low-rank approximation to enhance computation efficiency. (D) Output: For each gene, DenMark provides (1) posterior summaries of correlation between cell density and gene expression per cell; (2) the visualization of the latent field decomposition, where the gene-expression intensity is expressed as the weighted sum of a) a shared spatial component driven by the same latent field of cell density, and b) a residual component that reflects gene-specific spatial structure beyond what cell density accounts for; and (3) spatial correlation functions describing how correlations in cell density (or gene expression) decrease as a function of intercellular distance.

### Benchmark Evaluation of DenMark with Simulated Data

To verify that DenMark is a feasible method and characterize the density-expression correlation reasonably well, we perform simulation studies. One key step in DenMark is to use a low-rank approximation Gaussian process (i.e., HSGP) to achieve scalability for large datasets. At the same time, this efficiency comes at the cost of potential loss in the parameter estimation accuracy. Therefore, we first examine whether DenMark achieves scalability without sacrificing much in model performance. Here, we conduct a simulation study to compare the performance of the method employing the exact GP with that of the method whose implementation is facilitated by the HSGP approximation. In this simulation study, cell counts and aggregated gene expression levels are generated directly within each grid. In total, 100 synthetic datasets are generated, each simulated on a 22 × 22 designed grid within a square region of size 4 mm^2^. The details of the simulation are presented in Method. Figure 2A illustrates a typical simulated dataset, which shows one gene whose expression is positively associated with cell density.

**Figure 2.**
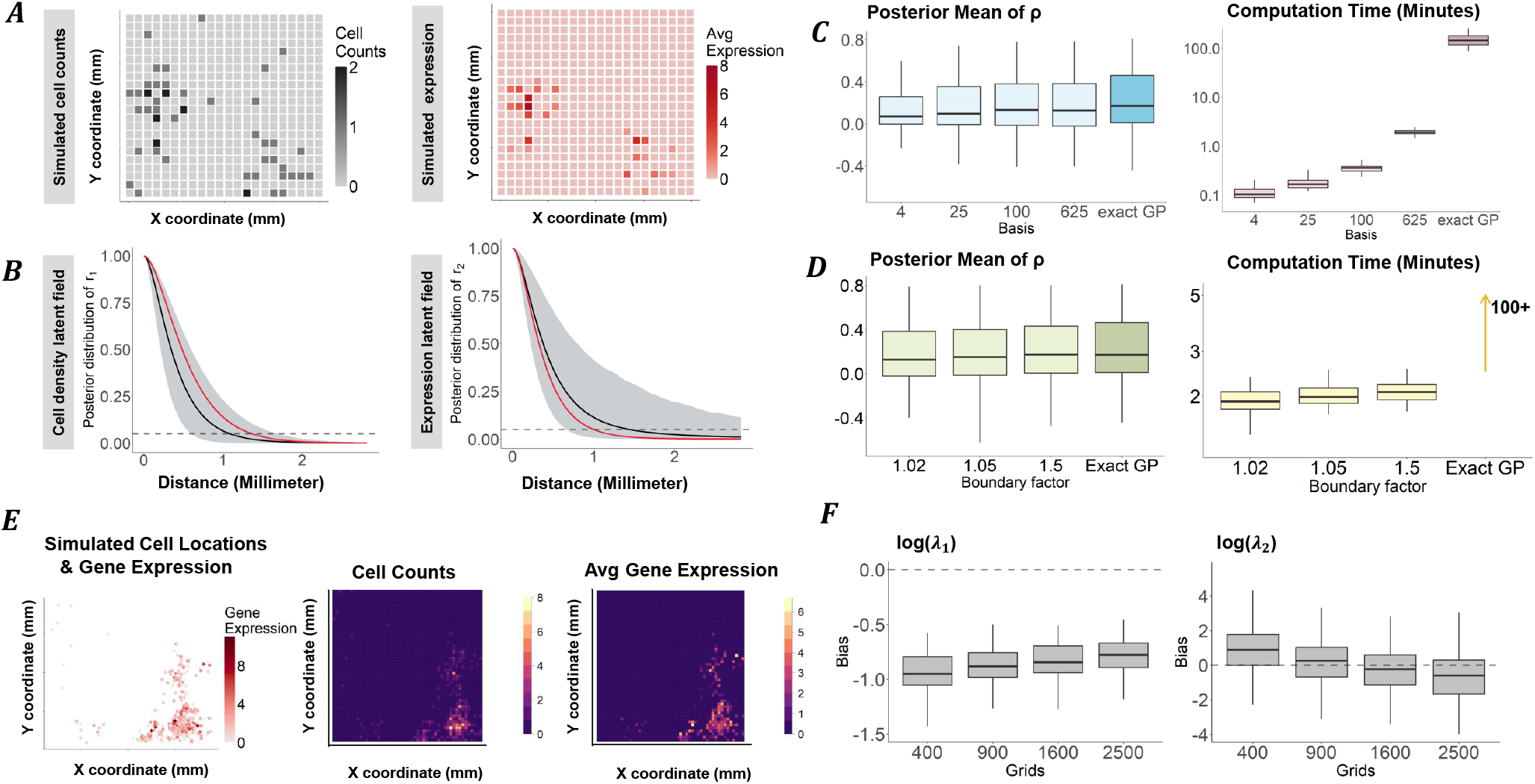
Results of the simulation studies. (A) Simulated cell counts and average gene expression per grid on 22 × 22 grids for an exemplary simulated dataset. (B) Spatial correlation functions of the latent spatial fields of cell density and gene expression, illustrating the decay of spatial correlation with increasing cell-cell distance (0 mm to the maximum observed distance). Black solid line, posterior mean; grey shaded region, 95% credible interval; red line, true correlation function. The horizontal dashed line (0.05) denotes a practical threshold below which correlation is considered negligible. (C) Comparison of the exact Gaussian process (GP) and Hilbert space Gaussian process (HSGP) approximations with different numbers of basis functions (2 × 2, 5 × 5, 10 × 10, and 25 × 25 over the two-dimensional spatial domain), together with corresponding computational time. (D) Comparison of exact GP and HSGP under varying boundary expansion factors (1.02, 1.05, and 1.5 applied to two spatial directions), with corresponding computational time. (E) Simulated marked point process (left), corresponding binned cell counts (middle), and average of gene expression per bin (right). (F) Estimation bias between fitted grid-level and simulated cell-level log-intensity surface across predefined grid resolutions (20 × 20, 30 × 30, 40 × 40 and 50 × 50).

In the analysis of the simulated data, we first fit the model with an exact Gaussian process approach as a reference for comparison. Spatial correlation functions are shown to assess how spatial correlation decays as a function of the Euclidean distance between cells (black solid lines; see Method), within the cell density (Figure 2B, left panel) and single-cell gene expression (Figure 2B, right panel), respectively. For benchmarking, we compare them to their true spatial correlation functions (Figure 2B, red lines). The 95% credible intervals (shaded areas) are shown to cover the true correlation functions in both spatial patterns, indicating DenMark accurately captures the true spatial correlation structure. We then evaluate the accuracy of the parameter estimates, which shows that the posterior means closely matched their true parameter values, indicating accurate estimation of the model parameters (Figure S1).

We then compare the exact GP to the approximations obtained using HSGP with various numbers of basis functions. In the implementation of HSGP, the number of basis functions determines the trade-off between approximation accuracy and computational efficiency, whereas the boundary factor, a domain-padding parameter in the HSGP basis, is essential in reducing edge effects (Riutort-Mayol et al., 2023). Therefore, we examine the posterior spatial correlation under various configurations of these two key hyperparameters. For the number of basis functions, we set it to be 4, 25, 100 and 625, respectively, to explore its influence on the performance. Here, the primary parameter of interest is the correlation between cell density and gene expression processes, i.e., *ρ* (see Method). Models with HSGP approximation performance achieved are observed to be comparable to those of the exact GP model, and their differences decrease as the number of basis functions increases (Figure 2C). On the other hand, computation time increased rapidly with the number of basis functions. These results confirm that a reasonably chosen approximation adopted in DenMark can achieve an effective balance between accuracy and efficiency. Then, we investigate boundary factors which are set to 1.02, 1.05, and 1.5, with the number of basis functions fixed as 625. Both the accuracy results and computation time increase only slightly with a larger boundary factor, as shown in Figure 2D, suggesting that DenMark’s performance is relatively robust to the choice of boundary factor at this specific choice of the number of basis function.

Because the corresponding likelihood function of DenMark involves an integral without a closed-form solution, numerical approximation is typically employed. We therefore approximate the single-cell-resolution point process and gene expression process using a grid-based discretization as in Figure 1B. Since this approach relies on a predefined gridding, the resulting inference may be influenced by how finely the grid is specified, and our second simulation study examines how sensitive the results are to the choice of grid size. Specifically, 100 datasets are simulated from a marked point process with known parameters as described in Method and the study domain is discretized by various grid levels, where the performance of the approximation bias are investigated. In an example, the simulated points together with their gene expressions are binned into 50 × 50 grids (Figure 2E). The model is then refit using the HSGP approximation, with the study domain discretized into grids of sizes 20 × 20, 30 × 30, 40 × 40, and 50 × 50, respectively, corresponding to different resolutions of the approximation. To evaluate the performance of the grid-based discretization, the simulated gridded cell counts and gene expression are treated as ground truth. They are then compared with the expected values from DenMark, and the grid-level bias is quantified as the difference between the estimate and the ground truth. The resulting comparison shows that as grid resolution increases, reconstructions of cell counts more closely match the gridded ground truth, and a similar trend is observed for gene expression (Figure 2F). These results suggest that when computational resources allow, a finer grid is preferable as it enhances the performance of grid-based discretization for the approximation.

### DenMark validates known and identifies novel density-correlated genes in the mouse brain

To illustrate how DenMark can help validate genes hypothesized to be associated with cell distribution, we analyze MERSCOPE mouse brain data from Vizgen, comprising coronal sections across three mouse brain tissue replicates (Vizgen, 2021). Our investigation focuses on one replicate (sample ID: S1R1), which includes 78,329 cell locations and 649 genes. After data quality control, we keep 483 genes at 78,329 cells. The cells are categorized into astrocyte, microglia, neuron (including both excitatory and inhibitory neurons) and other unannotated cells using cell-type marker genes (*Aqp4, Gfap*, and *Aldh1l1* for astrocyte cells; *P2ry12* and *Cx3cr1* for microglia cells; *Slc17a6, Slc17a7, Gad1* and *Slc32a1* for neurons) (Figure 3A). Since the region of interest (ROI) encompassed the entire tissue section, which contains a highly heterogeneous mixture of cell types, jointly modelling all cells, regardless of cell type, would likely obscure cell-type-specific spatial structure and density-expression relationships. To obtain stable and interpretable estimates of the latent intensity fields and the associated density-expression correlation, we model one cell type at a time and primarily focus on astrocytes. Astrocytes are among the most abundant cell types in this dataset (23,624 cells, Figure 3A), providing sufficient counts for reliable point process inference, and they exhibit clear large-scale spatial organization across the tissue. Moreover, given their central role in brain homeostasis and involvement in diverse neurological disorders (Schitine et al., 2015; Westergard and Rothstein, 2020), astrocytes offer a biologically meaningful context in which to illustrate the performance and interpretability of our framework. Within astrocytes, we focus on several genes with established links to cell adhesion, migration, and brain diseases: *Aqp4, Cxcl12, Cxcr4, Egfr*, and *Gfap. Aqp4* is a gene playing a key role in brain edema after injury and other brain diseases (Venero et al., 2001; Papadopoulos and Verkman, 2007), and has been implicated in astroglial cell migration (Saadoun et al., 2005; Ding et al., 2011). *Cxcl12* and its receptor encoding gene *Cxcr4* are essential in controlling cell migration and cell positioning (Quinn et al., 2018). One known pathway through which *Cxcl12* receptors promote cell signaling is through Src-mediated activation of the epidermal growth factor receptor, encoded by *Egfr* (Cheng et al., 2017; Zieger-Naumann et al., 2023). *Gfap* encodes glial fibrillary acidic protein (GFAP), a canonical intermediate filament of astrocytes that is upregulated during reactive astrogliosis, which is a process characterized by local changes in astrocyte density and morphology.

**Figure 3.**
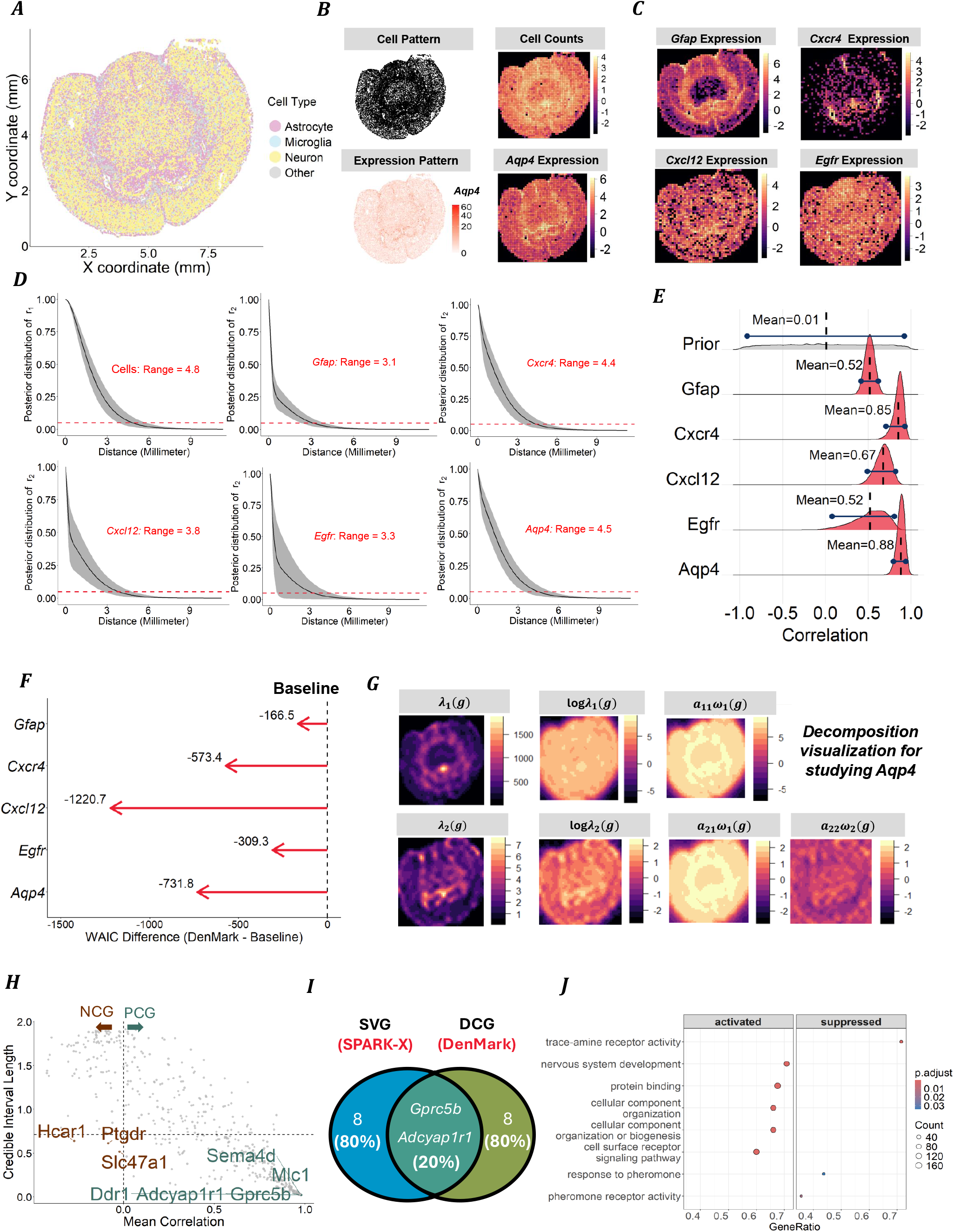
Results for the data analysis on Vizgen MERFISH Mouse brain. (A) Cell type annotation of the whole mouse brain *Slice 1, Replicate 1*. Each cell (one dot) is annotated with astrocyte, microglia, neurons, or others. (B) Spatial cell pattern for the individual astrocyte cells (top left) and their aggregated cell counts in grids in logarithm scale (top right) within the whole tissue section, together with the expression level of *Aqp4* of each cell visualized (bottom left) and their aggregated average expression per grid in logarithm scale (bottom right). (C) Log-transformed total gene expression per grid for *Egfr, Cxcr4, Cxcl12* and *Gfap*. (D) Spatial correlation functions (black curve) for cell occurrence and expression for gene *Aqp4, Egfr, Cxcr4, Cxcl12*, and *Gfap*. The shaded grey area indicates the 95% credible intervals. The red dashed horizontal line denotes the 0.05 correlation threshold, below which spatial dependence is considered negligible in practice; the distance to the curve’s intersection with this threshold defines the practical range. (E) Prior and posterior distribution of the correlation between the cell density and gene expression latent fields. A common LKJ(1.2) prior is specified for the density-expression parameter, and the corresponding posterior distributions are shown for candidate genes. (F) WAIC comparison of the full model (DenMark), which allows dependence between gene expression and cell density, and a baseline model assuming independence, across candidate genes *Aqp4, Egfr, Cxcr4, Cxcl12*, and *Gfap*. The x-axis represents the WAIC difference (full minus baseline model); lower WAIC values indicate a better model fit. (G) Spatial patterns and latent field decomposition of astrocyte cell density pattern and *Aqp4* expression following DenMark framework. The first column shows the observed cell density and *Aqp4* expression intensity across spatial grids. The second column presents the corresponding log-transformed values. The third and fourth columns show the latent field decompositions of cell density and gene expression intensity, respectively. (H) Scatter plot of gene-specific associations between cell density and gene expression. Each point represents a gene, with the posterior mean density-expression correlation shown on the x-axis and the corresponding credible-interval length on the y-axis. The vertical dashed line indicates zero correlation, separating negatively density-correlated genes from the positive ones. The horizontal dashed line marks the median credible-interval length as an uncertainty threshold. Genes below this line exhibit more informative association estimates. Genes with the largest-magnitude correlations and shortest credible intervals are highlighted. (I) Venn diagram comparing the top 10 DCGs identified by DenMark and the top 10 SVGs detected by SPARK-X. The intersection represents a gene exhibiting both spatially heterogeneous expression and significant correlation with cell density. (J) Rank-based gene set enrichment analysis (GSEA) of the identified DEGs. Each dot represents a significantly enriched biological process, with activated and suppressed pathways shown separately. The x-axis indicates the gene ratio, and the y-axis lists enriched biological processes. Dot color encodes the FDR-adjusted p-value (p.adjust), and dot size reflects the number of contributing genes.

Cell locations and gene expression patterns were discretized onto 50 × 50 grids for *Aqp4* (Figure 3B), *Cxcl12, Cxcr4, Egfr*, and *Gfap* (Figure 3C). For each gene, we apply DenMark to the gridded gene expression and the cell density. We first focus on the spatial correlation by examining spatial correlation functions (Figure 3D). Spatial correlations between two locations are observed for both density and gene expression, but they diminish to negligible levels when the distance between them exceeds 5 millimeters. In general, the spatial correlation of gene expression decays more rapidly than that of astrocyte cell density. This suggests that the astrocyte density varies smoothly across the tissue in comparison to the spatial variation of the gene expression. Second, we find that *Aqp4* and *Cxcl12* exhibit more gradual spatial decay in expression than *Egfr, Cxcl12*, and *Gfap*, indicating broader spatial variation in their expression patterns. Then, we focus on quantifying the relationship between density and expression, inferred from the primary parameter of DenMark, *ρ*. DenMark provides the posterior distribution of *ρ* evaluated at the five selected genes together with the distribution of the assigned LKJ prior for comparison (Figure 3E). All posterior means are shown to be positive, which indicates the local average expression of these genes per cell is positively correlated to density. Among these genes, *Aqp4, Cxcr4* and *Cxcl12* have exhibited relatively higher density-expression correlation. The density-expression parameters provide continuous measures of association between local average gene expression and local cell density and therefore do not directly indicate whether a given gene should be regarded as density-correlated. To obtain a binary recommendation for each gene, we compare a model with the density-expression term to a model without including it, using the Watanabe-Akaike information criterion (WAIC) (see Method for details). The comparison suggests that, for all 5 candidate genes, the WAIC differences are strongly negative (Figure 3F), indicating that incorporating the density-expression term improves model fit and supporting the conclusion that each gene is density-correlated. Then, to understand how gene expression is associated with cell density, we visualize the gene expression pattern decomposition relative to the cell density pattern, as in Figure 1D. The majority of the *expression of Aqp4* (Figure 3G) and *Cxcr4* (Figure S2) can be explained by the component shared by cell density, with a minor contribution from the gene-specific component. As a comparison, a greater proportion of gene-specific variation remains for *Cxcl12, Egfr*, and *Gfap* (Figure S2).

Motivated by the encouraging validation results for the selected genes, we extend our analysis to the remaining 478 genes that meet the pre-processing criteria and apply DenMark to each. For each gene, we examine the strength of the density-expression correlation as well as its uncertainty (Figure 3H). We identify that *Gprc5b, Adcyap1r1, Mlc1, Sema4d*, and *Ddr1* show the strongest positive correlation with cell density, while *Hca1, Ptgdr*, and *slc47a1* exhibit the most negative correlation with cell density. Importantly, DCGs are not necessarily strongly spatially varying, as they are conceptually distinguished. We further examine the overlap between the top 10 DCGs (see Data S1, provided as a CSV file in the Supplemental Information) and the top 10 SVGs identified by SPARK-X (Figure 3I). Only two genes (*Gprc5b* and *Adcyap1r1*) are shared, exhibiting both strong spatial variation and astrocyte-density correlation. Finally, to assess whether the identified DCGs are associated with known biological processes, we performed a rank-based gene set enrichment analysis (GSEA) using Gene Ontology (GO) terms for the selected DCGs. The identified DCGs are highly enriched for well-established biological processes, including protein binding, nervous system development, cellular component organization, and cell-surface receptor signaling, consistent with structural remodeling and enhanced signaling in dense astrocyte regions. In contrast, the identified genes are suppressed in the trace-amine receptor activity, suggesting that this chemosensory pathway is relatively more active in low-density regions. These results provide validation that the identified genes reflect the biological signals associated with the underlying spatial and density-correlated structure, rather than artifacts arising from random gene selection.

### DenMark reveals immune-cancer interactions via density-correlated genes in the breast tumor microenvironment

Having shown that DenMark can identify density-correlated genes (DCGs) in whole-section brain tissue, we next investigate whether the framework generalizes to a distinct biological context and data-generating platforms. We apply DenMark to 10x Xenium spatial transcriptomics data from human breast tumors (Janesick et al., 2023), focusing on local tumor microenvironments rather than the entire tissue section to identify DCGs that characterize the surrounding niche. Compared with the whole-section analysis, the cell-type composition within a local tumor microenvironment is simpler and more homogeneous, allowing us to investigate multiple cell types simultaneously within the same niche. Consequently, the biological interpretation of DenMark in this setting is expected to differ from that obtained in whole-section analyses. This dataset contains 313 genes detected on 2 consecutive slices from a breast cancer patient. Our analysis focuses on the first sample (ID: Replicate 1), which includes expressions of 313 genes from 167,780 cells. We focus on three regions of interest (ROI), reflecting different microenvironments, which are respectively dominated by invasive tumor, DCIS-1, and DCIS-2 cells (Janesick et al., 2023). The ROIs are chosen to be 1 *mm*^2^ windows following pathologic annotation in Janesick et al. (2023). After preprocessing, we keep 243 genes at 5031 cells (3.0% of all cells in the entire tissue) in the DCIS-1 ROI, 246 genes from 3812 cells (2.3%) in the DCIS-2 ROI, and 222 genes from 6645 cells (4.0%) in the invasive tumor ROI (Figure 4A).

**Figure 4.**
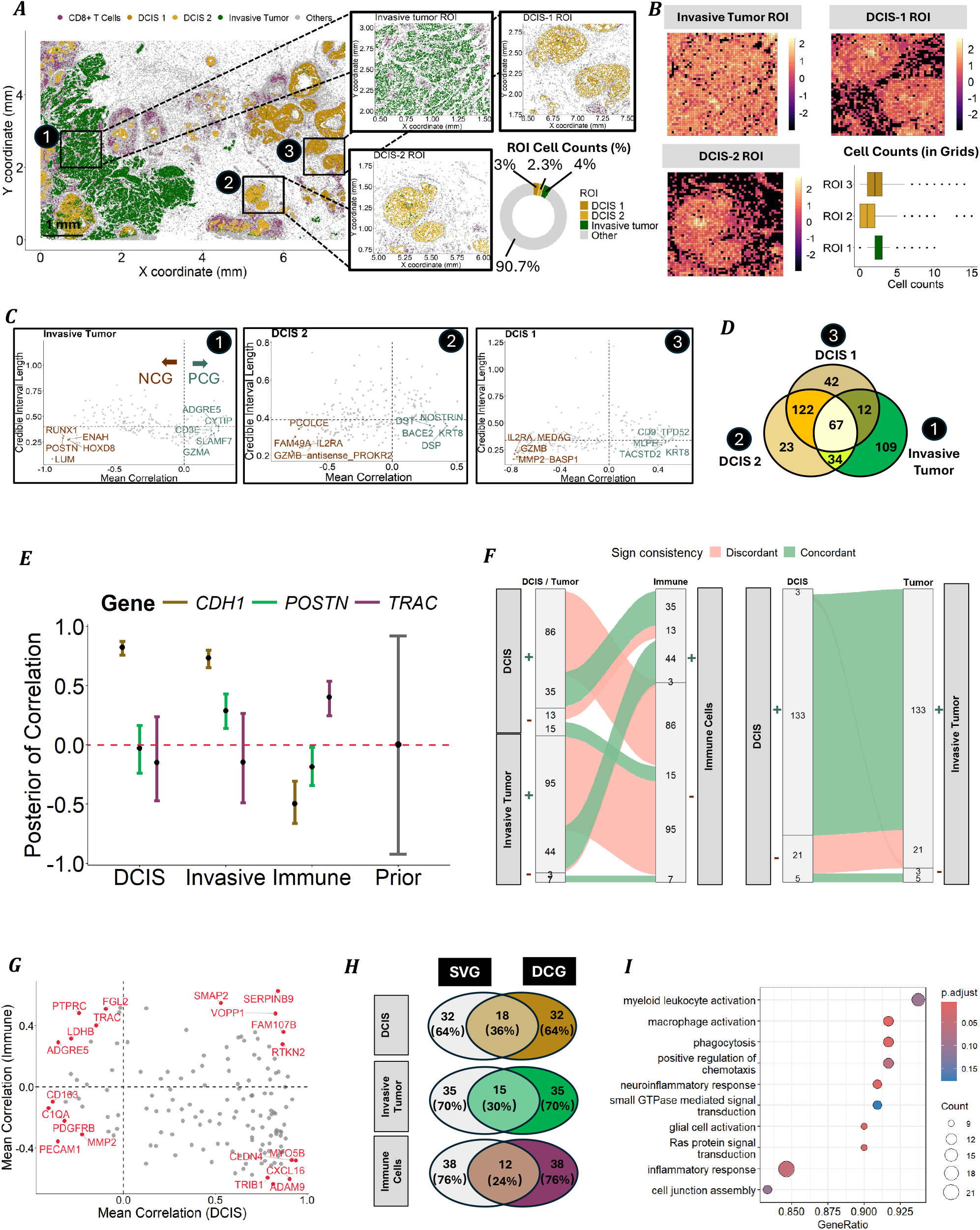
Microenvironment and whole tissue analysis on the 10x Xenium breast cancer. (A) Cell type annotation of the first breast cancer replicate with three zoomed-in, highlighted ROIs: pathological annotated Invasive tumor, DCIS-1, and DCIS-2 regions. The proportion of cells in each ROI relative to the full tissue section is summarized. (B) Total cell counts per grid for the invasive tumor ROI (top left), DCIS-1 (top right), and DCIS-2 (Bottom left) cells. Boxplot visualizing the distribution of total cell counts per grid in the three ROIs. (C) For each ROI (invasive tumor, DCIS2, and DCIS1), scatter plots show the posterior mean correlation between local cell density and gene expression (x-axis) and the corresponding credible-interval length (y-axis). The vertical dashed line marks zero correlation (negative density-correlated genes, NCG; positive density-correlated genes, PCG), and the horizontal dashed line indicates the uncertainty threshold; annotated genes show the strongest, most precisely estimated associations. (D) Venn diagram of density-correlated genes (DCGs) shared across ROIs. Overlap of DCGs identified in (1) invasive tumor, (2) DCIS2 and (3) DCIS1 regions. (E) Posterior density-expression correlations for pre-selected genes. Posterior mean correlation (points) with credible intervals (bars) for CDH1, POSTN, and TRAC in DCIS cells, invasive tumor cells, and CD8+ T cells, compared with the prior (right). The red dashed line indicates zero correlation. (F) Concordance of DenMark-identified density-correlated genes across cell populations. Sankey diagrams summarize the overlap of positively (+; PCG) and negatively (−; NCG) density-correlated genes between (left) DCIS, invasive tumor, and immune cells, and (right) DCIS tumor and invasive tumor cells. Green flows indicate concordant sign (PCG/PCG or NCG/NCG), whereas pink flows indicate discordant sign (PCG/NCG); numbers denote gene counts. (G) Comparing density-expression associations between DCIS and immune cells. Scatter plot of posterior mean correlations for each gene in DCIS (x-axis) versus immune cells (y-axis); dashed lines mark zero correlation in each population. The five genes with the largest absolute correlations in each quadrant are highlighted. (H) Overlap between DenMark DCGs and SPARK-X SVGs. Venn diagrams compare the top 50 density-correlated genes (DCGs; DenMark) with the top 50 spatially variable genes (SVGs; SPARK-X) in DCIS, invasive tumor, and CD8+ T cells; numbers (percentages) indicate overlap and method-specific genes. (I) Functional enrichment of DenMark-prioritized genes in invasive tumor cells. Dot plot of GO biological process enrichment based on genes ranked by DenMark, showing enriched immune and inflammatory pathways. The x-axis indicates GeneRatio, dot size denotes the number of genes (Count), and color encodes FDR-adjusted p-values (p.adjust).

After aggregating the cell counts in each ROI into 50 × 50 grids (Figure 4B), three ROIs exhibit distinct patterns of cell density variation over the region. Both DCIS-1 and DCIS-2 exhibit a markedly patchy density landscape with clear hotspots and sparse regions, whereas the invasive tumor ROI is comparatively more uniform, remaining denser overall. Given these distinct baseline density patterns, we analyzed each ROI separately. Within each ROI, DenMark yields a posterior distribution for the density-expression correlation for every gene, enabling us to classify genes as positively density-correlated or negatively density-correlated (Figure 4C). The resulting DCG sets are only partially shared across microenvironments (Figure 4D): DCIS-1 and DCIS-2 show the largest overlap, consistent with their shared in situ context, whereas the invasive tumor ROI contains the largest ROI-specific component. Notably, 67 DCGs are consistently identified in all three ROIs (see Data S2, provided as a CSV file in the Supplemental Information), suggesting a core set of genes whose expression robustly tracks local cellular patterns across distinct tumor niches.

Inspired by the partial overlap of DCGs across ROIs (Figure 4D), we next move beyond microenvironment-specific signals and perform a whole-section analysis to characterize global density-expression structure, with a focus on the spatial contrast between immune infiltration and malignant cell-dominated regions. After preprocessing, we retained 180 genes across 12923 cells for DCIS-1, 180 genes across 11683 cells for DCIS-2, 174 genes across 34374 cells for Invasive tumor cells, and 202 genes across 6940 cells for CD8+ T cells (Immune cells). We merged DCIS-1 and DCIS-2 into a single DCIS group, applied DenMark separately to DCIS, invasive tumor, and CD8+ T cells, and compared DCG profiles across populations.

To investigate antagonistic tumor-immune relationships, we examine three canonical marker genes spanning the major tissue compartments: *CDH1* (epithelial cell-cell adhesion, tumor-enriched), *POSTN* (secreted stromal remodeling program), and *TRAC* (T-cell marker enriched in immune regions) (Figure 4F). Consistent with compartment identity, *CDH1* shows positive posterior correlation in DCIS and invasive tumor but negative correlation in immune regions, whereas *TRAC* exhibits the opposite trend. *POSTN* displays a distinct domain-dependent pattern, highlighting microenvironmental remodeling across lesion and immune contexts. Model comparisons based on WAIC further support these gene- and cell-type-specific density-expression relationships (see section ‘WAIC criteria for model comparison’ in the Supplemental Information).

We then assessed whether DCGs exhibited consistent directions of density-expression correlation across populations (Figure 4E). We observed substantially more discordance between tumor cells and CD8+ T cells, whereas more DCGs are concordant between two tumor stages (DCIS vs invasive tumor; see Figure S4 in the Supplemental Information). This pattern suggests opposing density-expression signatures between malignant and immune compartments: transcripts that increase with tumor-cell density tend to decrease with immune-cell density, and vice versa. We find several genes strongly illustrate this antagonism, with immune markers (e.g., *PTPRC, TRAC*) opposing DCIS-enriched genes (e.g., *CLDN4, CXCL16*) (Figure 4G).

We compared the top 40 DCGs (see Data S2) with the top 40 SVGs (Figure 4H) identified by SPARK-X and observed only modest overlap across compartments (12-18 genes; 24-36%). This limited concordance indicates that DCGs capture a signal that is largely distinct from conventional spatial variability.

Rank-based GSEA in invasive tumor cells reveals enrichment of immune and myeloid-related biological processes among the DCG-ranked genes, including myeloid leukocyte activation, macrophage activation, phagocytosis, chemotaxis, and inflammatory response (Figure 4I). These enrichments are consistent with the established prominence and functional impact of tumor-associated myeloid populations in shaping the tumor microenvironment. In parallel, Ras protein signal transduction is enriched, implicating pathways linked to cell proliferation and migration and tumor invasion. Finally, terms annotated with glial cell activation and neuroinflammatory response likely reflect shared innate transcriptional programs rather than CNS-specific glial biology, pointing to strong myeloid-associated inflammatory signaling.

## DISCUSSION

In this paper, we propose DenMark, a model-based approach to quantify the spatial correlation between cell density and gene expression at single-cell resolution using ST data. DenMark incorporates a linear combination of GPs with a low-rank GP approximation, which preserves the spatial correlation between cell density and gene expression in GP while reducing computation time, for cell density and gene expression separately as a scalable framework. This model structure enables the estimation of the density-expression correlation. We show the application of DenMark on two single-cell level ST datasets (MERFISH mouse brain and Xenium breast cancer datasets), quantifying the density-expression correlation in the mouse brain and tumor-immune microenvironment. Additionally, DenMark enables the discovery of density-correlated marker genes in single-cell-resolution ST data. Using GSEA, we identified that these DCGs are correlated with activation of the nervous system and protein binding, and with suppression of trace-amine receptor activity in mouse brain ST data. In the invasive breast cancer environment, these identified DCGs are associated with cell junction assembly, activation of protein-DA complexes, and suppression of external encapsulation and extracellular matrix.

DenMark assumes spatially invariant elements in the coregionalization matrix, which may be restrictive when cell density or gene expression varies strongly across the ROI. Spatially varying coregionalization models could relax this assumption. DenMark also analyzes a single gene in one cell type, simplifying inference but ignoring gene-gene interactions. Extensions to multiple genes, slices, or cell types are possible. Finally, PCGs and NCGs are defined using the median credible interval of the posterior density-expression correlation. Alternatives, such as WAIC comparisons, could be considered.

## DATA AND CODE AVAILABILITY

(1) Mouse brain MERSCOPE data [https://console.cloud.google.com/storage/browser/public-datasets-vizgen-merfish]; (2) 10x Xenium data: [https://www.10xgenomics.com/products/xenium-in-situ/preview-dataset-human-breast] are publicly available as of the date of publication. All other data supporting the findings of this study are available within the article and its supplementary files. All other data reported in this paper will be shared by the lead contact upon request. An open-source implementation of the DenMark framework can be downloaded from https://github.com/StaGill/DenMark.

## ACKNOWLEDGEMENTS

This research was supported by the Natural Science and Engineering Research Council of Canada (NSERC) and the Canadian Statistical Science Institute (CANSSI). Zhang is a Fonds de recherche du Québec Research Scholar (Junior 1). His research was undertaken, in part, thanks to funding from the FRQ-Santé Program.

## AUTHOR CONTRIBUTIONS

Conceptualization, A.S and Q.Z.; methodology, A.S, M.X., and Q.Z.; investigation, M.X.; writing-original draft, M.X., A.S., and Q.Z.; writing-–review & editing, A.S. and Q.Z.; funding acquisition, A.S. and Q.Z.; supervision, A.S. and Q.Z.

## SUPPLEMENTAL INFORMATION INDEX

Figures S1-S4, and Table S1-S2 in a PDF

Data S1. List of DCGs identified in the mouse brain MERFISH dataset

Data S2. List of DCGs identified in the human breast cancer 10x Xenium dataset

## METHODS

### Method details

#### Region of Interest and Coordinate System Normalization

We consider a region of interest (ROI) within a tissue slice *D*_1_ ⊂ ℝ^2^, defined in a two-dimensional Cartesian coordinate system, as the spatial observation window. Our objective is to characterize the spatial distribution of cell occurrence and candidate gene expression within the specified ROI and to quantify their potential association. The choice of ROI is therefore critical and should balance sufficient cell counts with informative spatial variation in cell density. A smaller ROI is typically appropriate when the focus is on local microenvironmental structure, whereas a larger ROI is more suitable for investigating broader spatial patterns, where a large amount of cell counts support inference at larger spatial scales(Illian et al., 2008).

Suppose that *N* cells are observed within the tissue, whose locations on the tissue are denoted by {***s***_1_, ***s***_2_, · · ·, ***s***_*N*_} ⊂ *D*_1_, where each location ***s***_*i*_ = (*x*_*i*_, *y*_*i*_)^⊺^ ∈ ℝ^2^. Each observed cell *i* is associated with an RNA transcript count (called the mark) *m*(***s***_*i*_) ∈ *M* ⊂ 𝕄, which represents the gene expression measured at the location *s*_*i*_. Let ***m***_*N*_ = (*m*(***s***_1_), *m*(***s***_2_), · · ·, *m*(***s***_*N*_))^⊺^ denote the vector of gene expression measurements for the examined gene. Together, cell locations and their associated gene expression values constitute a marked point process (MPP), *N*_*m*_ = {[***s***_*i*_; *m*(***s***_*i*_)]}_*i*=1,2,···,*N*_, on the product space ℝ^2^ × 𝕄, where ℝ^2^ represents the spatial domain and 𝕄 denotes the mark space (Illian et al., 2008).

Our analysis begins with the observed cell spatial coordinates and gene expression measurements within the ROI. Spatial inference depends on the underlying coordinate reference system (CRS), so a unified, normalized definition of the ROI is required. To this end, cell coordinates {***s***_1_, ***s***_2_, · · ·, ***s***_*N*_} are rescaled to a common *mm × mm* coordinate system when necessary. In addition, the Hilbert space Gaussian process (HSGP) approximation, described in the following section, requires the spatial domain to be symmetric about the origin for the approximation to be well defined. The rescaled coordinates are therefore further transformed to a symmetric, bounded ROI (Roy and Schmidt, 2024), defined as *W* = [−*L*_1_, *L*_1_] × [−*L*_2_, *L*_2_]. The half ranges *L*_1_ and *L*_2_ are calculated by

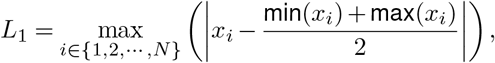

and

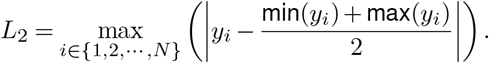

#### Data Preprocessing

For the gene expression data, we select genes based on two criteria: (1) variance across cells ranking in the top 95%, and (2) non-zero expression at no less than 5% of spatial locations. After filtering, *J* genes are kept and organized into a cell-by-gene expression matrix 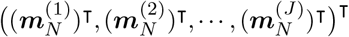, which summarizes the expression levels of *J* genes across the *N* observed cells within the ROI. Following quality control to remove non-informative genes, subsequent analyses focus on the remaining candidate genes. Each gene is analyzed independently, with one gene expression modeled at a time in relation to the spatial cells pattern within the ROI.

#### Density-dependent Marked Log Gaussian Cox Process

Our approach is built on a density-dependent marked point process, in which the spatial distribution of cells is used to characterize local cell density across the tissue. Gene-expression levels, treated as marks attached to individual cells, are modeled as count data, with expected expression reflecting two biological sources of variation: a shared component associated with the surrounding cellular environment (as summarized by local cell density) and a gene-specific component capturing expression patterns unique to each gene. This formulation allows the framework to describe tissue spatial organization and gene-expression heterogeneity, while quantifying how gene expression varies with changes in the local cellular microenvironment. As cell density is expected to vary inhomogeneously across a tissue slice and to exhibit stochastic spatial structure, we model the density-dependent marked point process using a Cox process framework (Banerjee et al., 2003). In particular, when the logarithm of the underlying cell density is modeled as a spatial Gaussian process, and gene expression is treated as a mark, the resulting formulation corresponds to a marked log-Gaussian Cox process (Møller et al., 1998; Ho and Stoyan, 2008; Myllymäki and Penttinen, 2009), as illustrated in fig. 1C. While this framework provides a flexible and biologically meaningful representation of spatial cellular organization and gene expression, its practical implementation is challenging because the integral term in the model yields intractable likelihoods (Møller et al., 1998). To address this issue, we approximate the integral using a strategy commonly referred to as the grid-based approach in the literature (Illian et al., 2013; Yadav et al., 2023).

#### Grid-based Approximation, Prior Specification and Spatial Correlation Function

As the grid-based approach relies on discrete Gaussian random fields, the underlying continuous spatial field is approximated on a discretized rectangular grid. These grids are employed to approximate the integral in the density-dependent marked log-Gaussian Cox process. Specifically, our framework partitions the rectangular region of interest *W* = [−*L*_1_, *L*_1_] × [−*L*_2_, *L*_2_] into *K* = *n*_row_ *× n*_col_ equally sized rectangular grids *G*_1_, *G*_2_, · · ·, *G*_*K*_ with a common area |*G*_1_| = · · · = |*G*_*K*_| = |*G*|, and let *g*_*i*_ denote the spatial location (centroid) of the grid *G*_*i*_. The continuous random field is therefore represented by a discretized counterpart, where the field is assumed to be constant within each grid. While finer grids yield more accurate approximations, they also add greater computational cost. To balance approximation accuracy with computational efficiency, the framework adopts a default resolution of *n*_*row*_ = *n*_*col*_ = 50, with finer resolutions available when higher precision is required. Within each grid, gene expression values from all contained cells are aggregated and normalized by the number of cells in that grid, yielding the cell count per grid recorded.

For each grid *G*_*i*_, let *N*_*i*_ denote the number of cells observed within that grid, and let *M*_*i*_ denote the aggregated gene expression obtained by summing expression values across all cells in the grid. Under the Cox process assumption, the cell count *N*_*i*_ in grid *G*_*i*_ is modeled as

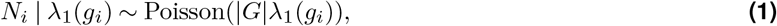

where *λ*_1_(*g*_*i*_) is the local cell density evaluated at the spatial location *g*_*i*_ corresponding to grid *G*_*i*_. The logarithm of the cell density is modeled as

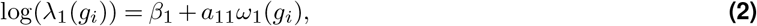

where *ω*_1_(***g***) is a Gaussian process with zero mean unit-variance with Matérn covariance (smoothness parameter fixed at 3/2) evaluated at the centroid vector of the grid ***g*** = (*g*_1_, · · ·, *g*_*K*_)^⊺^. This term represents a latent spatial effect that captures variation in cell density across the ROI. Under this discretization, the integral term in the density-dependent marked point process likelihood is approximated by a finite sum over the grids.

Similarly, single-cell gene-expression patterns are approximated at the grid level via the aggregated expression *M*_*i*_. Conditional on the observed cell count *N*_*i*_, gene expression in grid *G*_*i*_ is modeled as

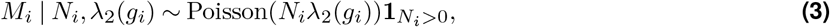

where the indicator 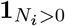 restricts gene expression to grids containing at least one observed cell. The gene-expression intensity *λ*_2_(*g*_*i*_) is modeled on the log scale as

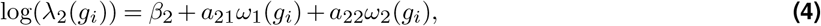

where *ω*_2_(***g***) is an independent zero-mean unit-variance Gaussian process with Matérn covariance (smoothness parameter fixed at 3/2) independent of *ω*_1_(***g***). The term *a*_21_*ω*_1_(*g*_*i*_) represents a shared spatial effect linking gene expression to local cell density, whereas *a*_22_*ω*_2_(*g*_*i*_) captures gene-specific spatial variation beyond that explained by cell density.

To facilitate prior specification and enhance interpretability, we express the latent field vector (*a*_11_*ω*_1_(*g*_*i*_), *a*_21_*ω*_1_(*g*_*i*_) + *a*_22_*ω*_2_(*g*_*i*_))^⊺^, corresponding to the grid-level cell density and gene expression latent fields, in matrix form:

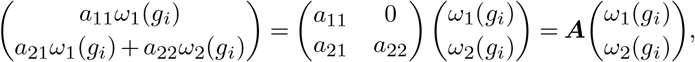

where ***A*** is the coregionalization matrix that governs how the latent Gaussian processes *ω*_1_(***g***) and *ω*_2_(***g***) jointly contribute to the spatial variation in cell density and gene expression. The marginal covariance between the log cell density and log gene expression latent fields at the same grid is given by ***AA***^⊺^. Following the covariance separation strategy (Barnard et al., 2000), we decompose this covariance into marginal standard deviations and a correlation structure,

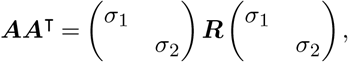

where *σ*_1_ and *σ*_2_ represent the marginal spatial variability of cell density and gene expression, respectively, and 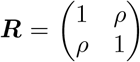 is the latent field correlation matrix. This covariance parametrization in terms of marginal standard deviations (*σ*_1_, *σ*_2_) and the density-expression correlation parameter *ρ* provides a biologically interpretable representation of shared and gene-specific spatial effects.

Under this parametrization, the correlation between the cell density log(*λ*_1_(*g*_*i*_)) and the gene expression intensity log(*λ* (*g*)) within grid *g* is given by 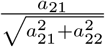 (see section ‘The derivation of the correlation between latent field decomposition in DenMark’ in the Supplemental Information). The magnitude of this correlation, 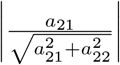 quantifies the strength of association between local cell density and gene expression while the sign of *a*_21_ determines its direction. This allows genes to be naturally classified as negatively density correlated (*a*_21_ *<* 0), density uncorrelated (*a*_21_ = 0), or positively density correlated (*a*_21_ *>* 0).

We next describe the prior distributions assigned to the model parameters Θ = (*β*_1_, *β*_2_, *σ*_1_, *σ*_2_, *ρ, ϕ*_1_, *ϕ*_2_)^⊺^ in the framework. Assuming prior independence across parameters, the joint prior distribution factorizes as

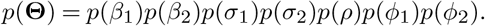

We adopt weakly informative priors for all parameters to stabilize inference while retaining sufficient flexibility to capture biologically meaningful variation. Specifically, we assign an LKJ(1.2) prior (Lewandowski et al., 2009) to the matrix ***R***, inducing an approximate uniform prior on the correlation parameter *ρ* that governs the dependence between the latent fields associated with cell-density and gene-expression. Intercepts are assigned independent normal priors with large variance, *β*_*l*_ ∼ *N* (0, 5^2^) for *l* = 1, 2. The marginal standard deviations of the latent fields are given half-normal priors, *σ*_*l*_ ∼ *N* ^+^(0, 1) for *l* = 1, 2, ensuring positivity while mildly constraining their scale. For the length-scale parameters *ϕ*_1_ and *ϕ*_2_ of the two Gaussian processes, we assign inverse-gamma priors *ϕ*_*l*_∼ InvGamma 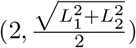. The scale parameter corresponds to half of the maximum spatial extent of the ROI, thereby encoding weak prior information on the rate at which spatial correlations decay and providing biologically reasonable guidance on the smoothness of the latent spatial processes.

Following Bayes’ theorem, the kernel of the posterior is given by

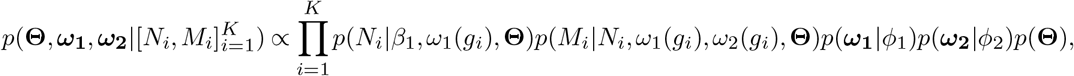

which does not have a closed-form solution. Therefore, we employ Hamiltonian Monte Carlo to sample from the posterior distribution.

We refer to Equations 1 and 3 with intensity structure Equations 2 and 4 as the <u>Den</u>sity-dependent <u>Mark</u>ed Point Process framework (**DenMark**). Under DenMark, spatial dependence in the Gaussian processes *ω*_1_(***g***) and *ω*_2_(***g***) is specified through correlation functions *ρ*_*k*_(*d*) = *ρ*_*k*_(|*g*_*i*_ − *g*_*j*_|), where *g*_*i*_, *g*_*j*_ ∈ {*g*_1_, · · ·, *g*_*K*_} denote spatial grid locations, and *d* is their Euclidean distance (*k* = 1, 2). Both correlation functions are specified using a Matérn covariance with smoothness parameter fixed at 3*/*2, where *ϕ*_1_ and *ϕ*_2_ denote the length-scale parameters governing the spatial decay of two Gaussian processes *ω*_1_(***g***) and *ω*_2_(***g***), respectively. This value of the smoothness parameter guarantees that the resultant processes are once mean-square differentiable.

To facilitate biological interpretation, we characterize spatial dependence using practical range parameters *r*_1_ and *r*_2_, which represent the distances beyond which spatial correlations in the cell-density-associated and gene-expression-associated latent fields, respectively, become negligible. We define the practical range as the distance at which the correlation function decays to 0.05, a commonly used threshold for spatial independence (Gelfand et al., 2004). Correspondingly, the practical range for the cell density latent field is obtained by solving

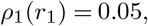

whereas the practical range for the gene-expression latent field is determined by solving

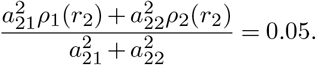

The estimated range parameters *r*_1_ and *r*_2_ provide intuitive measures of the spatial scales over which cell density and gene expression exhibit meaningful spatial correlation, and the corresponding spatial correlation functions can be visualized to aid interpretation.

#### Hilbert Space Gaussian Process Approximation

The grid-based approach requires evaluating Gaussian processes at a large number of spatial locations, which becomes computationally expensive as grid resolution increases. To address this limitation, DenMark incorporates an additional approximation based on a low-rank Hilbert space Gaussian process representation to efficiently approximate the GPs *ω*_1_(***g***) and *ω*_2_(***g***). Under this representation, the Gaussian process kernel *k*(·,·) for two grids *G*_*u*_ and *G*_*u*_*′* is approximated as

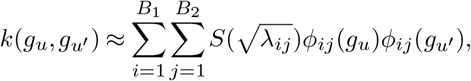

where *B*_1_ and *B*_2_ denote the prespecified numbers of basis functions along the x- and y-coordinates, respectively, *S*(·) is the spectral density function, and *λ*_*ij*_, *ϕ*_*ij*_(·) are the eigenvalues and eigenfunctions of the Laplacian operator on the spatial domain, respectively.

Correspondingly, each Gaussian process can be approximated as

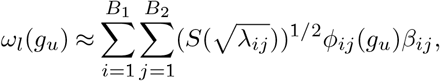

for *l* = 1, 2, where *β*_*ij*_ ∼ *N* (0, 1). This low-rank representation substantially reduces computational complexity while retaining the essential spatial structure of the original Gaussian processes.

Posterior draws for the model parameters Θ = (*β*_1_, *β*_2_, *σ*_1_, *σ*_2_, *ϕ*_1_, *ϕ*_2_, *ρ*)^⊺^ are performed using Hamiltonian Monte Carlo sampling, as implemented in Rstan.

#### Model Comparison: DenMark and Baseline Variant

For the coregionalization matrix ***A***, DenMark introduces a shared parameter *a*_21_ that links the cell density and gene expression, and quantifies the extent to which gene expression shares spatial structure with local cell density. To assess whether gene expression is associated with cell density, we further introduce a baseline density-independent version of DenMark in which gene expression varies independently of cell density. Specifically, we compare the full model to a nested specification obtained by constraining *a*_21_ = 0, as summarized in Table 1. Model comparison is conducted using the Watanabe-Akaike information criterion (WAIC), where small values indicate a more preferable model (see Supplemental Information, section ‘WAIC criteria for model comparison’ for details).

**Table 1.**
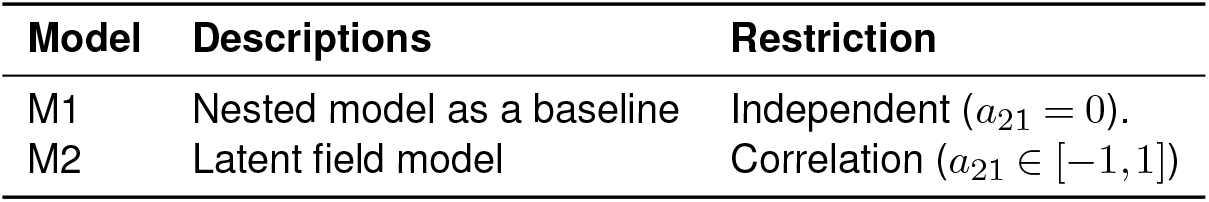
Model comparison between DenMark and a nested setting.

We further note that the DenMark (M2) specification generalizes the marked point process model of Illian et al. (2013) (Illian et al., 2013) by incorporating an additional gene-specific spatial component, *a*_22_*ω*_2_(***g***), in addition to the shared density-associated component. This extension explicitly captures gene expression patterns that are only partially explained by cell density, thereby improving biological interpretability and modeling flexibility.

