## Supplementary Information for "DenMark: A Bayesian Hierarchical Model for Identifying Cell-Density Correlated Genes from Spatial Transcriptomics"

### Simulation: Comparison between Exact GP and HSGP

In Figure 2 $A$, $B$, $C$, and $D$, we simulate spatial cell density and gene expression fields using a GP and compare inference obtained from the exact GP with that from the HSGP approximation. The true parameter values are set to be $\sigma_1 = 1$, $\sigma_2 = 1$, $\rho = 0.5$, $\beta_0 = 2$, $\beta_1 = 1$, $\phi_1 = 0.5$ and $\phi_2 = 0.25$. These values are chosen to ensure that the simulated cell counts and gene expression levels remain within biologically plausible ranges.

We first generate two independent latent spatial fields, $\omega_1(\boldsymbol{g})$ and $\omega_2(\boldsymbol{g})$, as zero-mean, unit-variance Gaussian processes with a Matérn covariance kernel with $\nu = 3/2$, as specified in Equations 1 and 3. Conditional on $\omega_1(g_i)$, the grid-level cell counts $N_i$ are simulated from a Poisson distribution,

$$N_i|\omega_1(g_i) \sim \text{Poisson}(|G|\exp(\beta_0 + a_{11}\omega_1(g_i))),$$

Given the observed cell counts, grid-level gene expression $M_i$ is generated as

$$M_i|N_i,\omega_1(g_i),\omega_2(g_i) \sim \begin{cases} 0 & \text{if } N_i = 0 \\ \text{Poisson}\Big(N_i \exp(\beta_1 + a_{21}\omega_1(g_i) + a_{22}\omega_2(g_i))\Big) & \text{if } N_i > 0. \end{cases}$$

The resulting simulated dataset consists of paired grid-level observations $\{[N_i, M_i]\}_{i=1}^{K}$.

We first fit the model using the exact GP formulation. We then refit the same model using the low-rank Hilbert space Gaussian process (HSGP) approximation, in which the number of basis functions and the boundary factor are the primary tuning parameters controlling approximation accuracy. Increasing either the number of basis functions or the boundary factor improves approximation accuracy at the cost of increased computational time. Based on this trade-off, we select a configuration of $25 \times 25$ basis functions with a boundary factor of 1.5, which balances accuracy and computational efficiency.

To further assess the sensitivity of the approximation, we examine the effects of these two tuning parameters separately. Holding the boundary factor fixed at 1.5, we vary the total number of basis functions across $4, 25, 100, 625$, with equal numbers assigned to the x- and y- axis. These values are chosen to inspect a range of approximation resolutions while maintaining numerical stability. Conversely, with the number of basis functions fixed at a sufficiently large value (625), we investigate the impact of different choices of boundary factors.

#### Simulation: Single-cell Resolution MPP Fitted with Grid-Cell Resolution Approximation

In fig. 2 $E$ and $F$, we simulate a marked point process to evaluate the accuracy of the grid-based approximation used to fit the density-dependent marked point process model. This simulation assesses how well the gridded likelihood recovers the underlying continuous latent cell-density and gene-expression-intensity fields when inference is performed on discretized spatial domains.

We simulate cell locations (points) and their associated gene-expression (marks) within a two-dimensional tissue window $[-1 \text{ mm}, 1 \text{ mm}] \times [-1 \text{ mm}, 1 \text{ mm}]$. Model parameters are set to $\beta_0 = 6.2$, $\beta_1 = -1$, $a_{21} = 0.6$, $\sigma_1 = 2$, $\sigma_2 = 1.1$, $\phi_1 = 0.8$ and $\phi_2 = 0.4$. These values are informed by the pilot analysis of mouse brain MERFISH data and further calibrated to yield biologically plausible ranges of local cell density and per-cell gene expression intensity. Let $\mathbf{\Lambda}_1(s) = (\Lambda_1(s_1), \cdots, \Lambda_1(s_N))^\intercal$ denote the true underlying spatial intensity of cells, and $\mathbf{\Lambda}_2(s) = (\Lambda_2(s_1), \cdots, \Lambda_2(s_N))^\intercal$ denote the true gene expression intensity evaluated at the simulated cell locations.

We apply grid-based aggregation to approximate the continuous latent field using $50 \times 50$ grids. This discretization implicitly defines spatial grids within which cells are assumed to experience similar local tissue microenvironments. The model is then fitted to the resulting grid-level summaries, yielding posterior mean estimates of the latent cell density and gene expression intensities, $\boldsymbol{\lambda}_1(g) = (\lambda_1(g_1), \cdots, \lambda_1(g_{2500}))^\intercal$ and $\boldsymbol{\lambda}_2(g) = (\lambda_2(g_1), \cdots, \lambda_2(g_{2500}))^\intercal$. To compare the fitted grid-based model with the underlying continuous biological process, we assign each simulated cell location $s$ the posterior mean intensity of the grid containing it, $\widehat{\lambda_1}(s) = \lambda_1(g_i)$, and $\widehat{\lambda_2}(s) = \lambda_2(g_i)$, for $s \in g_i$. This step reflects the biological assumption that all cells within a small spatial neighborhood share a common microenvironmental signal.

We quantify the discrepancy between the estimated grid-level intensities (cell density and gene expression intensity) and the true point-level intensities by computing the average bias on the log scale. Specifically, the bias in estimated cell density is defined as

$$\text{Bias}_{\log \lambda_1} = \frac{1}{N} \sum_{s \in \{1,2,\ldots,N\}} \left( \log(\widehat{\lambda}_1(s)) - \log(\lambda_1(s)) \right),$$

and the corresponding bias in gene expression intensity is

$$\text{Bias}_{\log \lambda_2} = \frac{1}{N} \sum_{s \in \{1,2,\ldots,N\}} \left( \log(\widehat{\lambda}_2(s)) - \log(\lambda_2(s)) \right).$$

These bias measures capture how well the grid-based model preserves underlying biological variations in cell density and gene expression as using the grid-based approximation.

607 # Simulation Design: Parameter Recovery

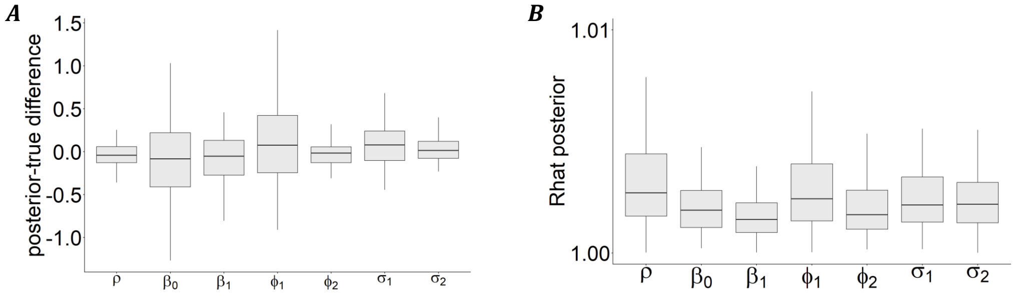


**Figure S1.** Posterior recovery of model parameters. (A) Differences between posterior estimates of each parameter and the corresponding true value. (B) Posterior distributions of the $\hat{R}$ convergence statistic for each model parameter.

### Analysis on the Mouse Brain

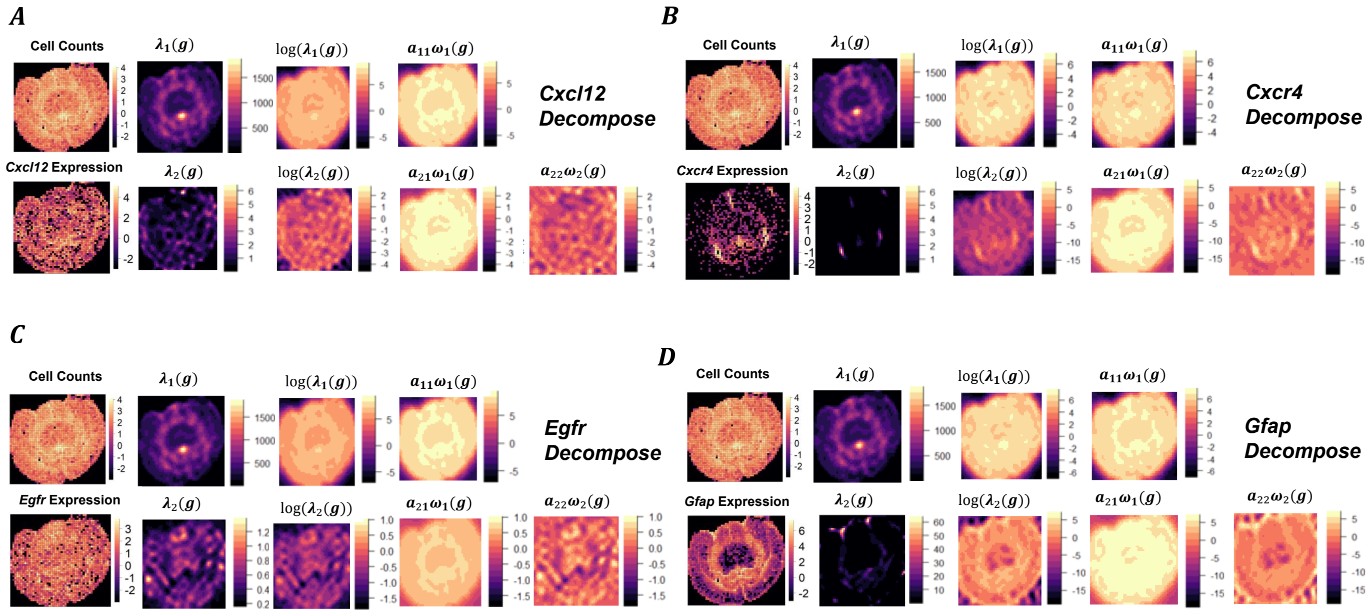


**Figure S2.** Latent field decomposition for genes *Cxcl12*, *Cxcr4*, *Egfr* and *Gfap*.

#### Analysis on the 10X Xenium Breast Cancer

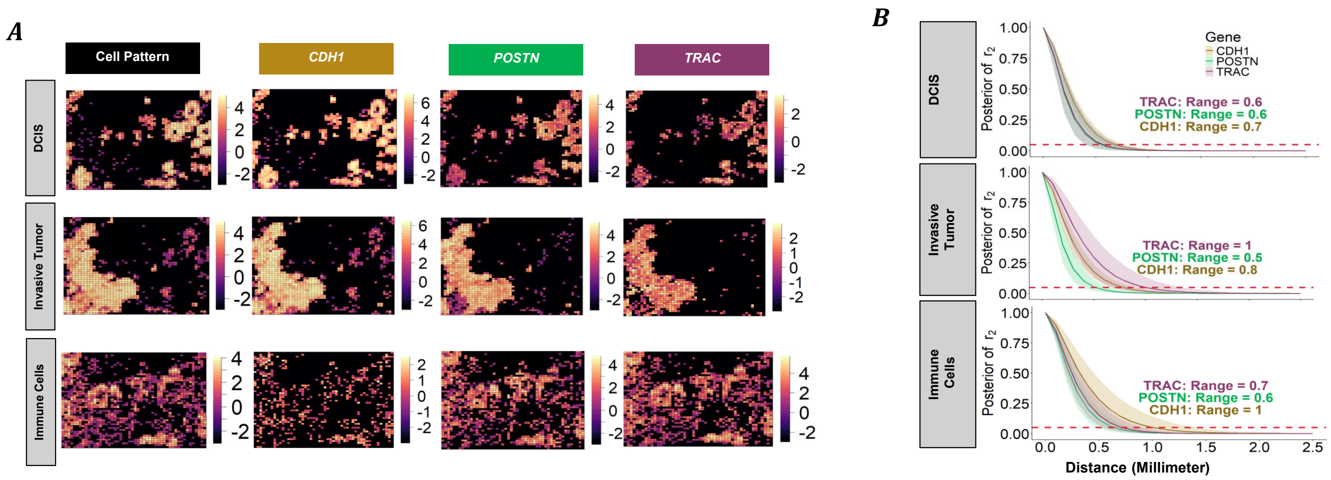


**Figure S3.**
(A) Cell pattern versus the *CDH1*, *POSTN*, and *TRAC* gene expressions at DCIS, invasive tumor, and immune cells. Cell counts and gene expression at each grid are scaled in a logarithmic scale.
(B) Spatial Correlation functions from the whole tissue analysis for genes *CDH1*, *POSTN*, and *TRAC* at DCIS cells, invasive tumor cells, and CD8+ T immune cells, respectively.

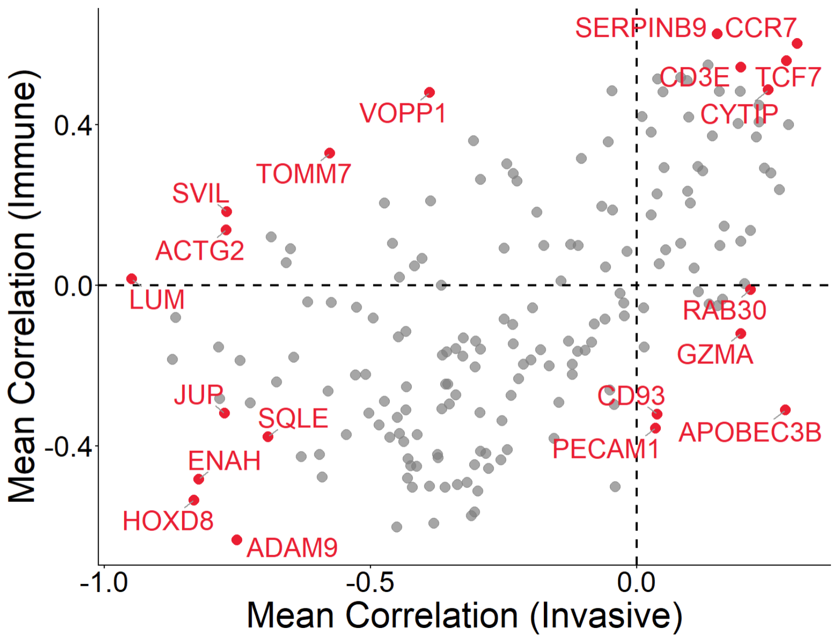


**Figure S4.** Analysis on the whole tissue on the Invasive & CD8+ immune cells. Dots represent the posterior mean of the tumor cells and immune cells. Five genes with the largest distance to the origin (0,0) in each of the four quadrants are highlighted in red.

### Derivation of the Correlation between Latent Field Decomposition in DenMark

Correlation in the DenMark with Equations 1 and 3 with intensity structure Equations 2 and 4 can be calculated by

$$\begin{aligned}
\text{Corr}(\beta_1 + a_{11}\omega_1(s), \beta_2 + a_{21}\omega_1 + a_{22}\omega_2(s)) &= \frac{\text{Cov}(\beta_1 + a_{11}\omega_1(s), \beta_2 + a_{21}\omega_1(s) + a_{22}\omega_2(s))}{\sqrt{\text{Var}(\beta_1 + a_{11}\omega_1(s))}\sqrt{\text{Var}(\beta_2 + a_{21}\omega_1 + a_{22}\omega_2(s)))}} \\
&= \frac{a_{11}a_{21}}{\sqrt{a_{11}^2}\sqrt{a_{21}^2 + a_{22}^2}} \\
&= \frac{a_{21}}{\sqrt{a_{21}^2 + a_{22}^2}}.
\end{aligned}$$

#### WAIC Criteria for Model Comparison

Model comparison is designed to assess whether a shared density-dependent part ($a_{21}\omega_1(\mathbf{g})$) is necessary, and the comparison between DenMark and the benchmark is based on WAIC computed for each model.
Let the observed data be $\mathcal{D} = \{(N_i, M_i) : i = 1, 2, \cdots, K\}$. Let $\Theta = (\beta_1, \beta_2, \sigma_1, \sigma_2, \rho, \phi_1, \phi_2)^\mathsf{T}$ denote all unknown parameters and latent effects. We compare DenMark with the baseline model in Watanabe-Akaike information criterion (WAIC) criteria. For DenMark, the WAIC is calculated by

$$\begin{aligned}
\text{lpdd} &= \sum_{i=1}^{K} \log\left(\frac{1}{Q-M} \sum_{m=M+1}^{Q} p(N_i, M_i | \boldsymbol{\Theta}^{[m]})\right), \\
\text{pwaic} &= \sum_{i=1}^{K} \text{Var}_{m=M+1}^{Q} \log(p(N_i, M_i | \boldsymbol{\Theta}^{[m]})), \\
\text{WAIC} &= -2(\text{lpdd} - \text{pwaic}),
\end{aligned}$$

where the superscript $[m]$ denotes a draw from the posterior distribution of the parameter, $M$ is the size of the burn-in period, $Q$ is MCMC sample size, and $\text{Var}_{m=M+1}^{Q} z_m = \frac{1}{Q-M} \sum_{m=M+1}^{Q} (z_m - \bar{z})^2$ represents the sample variance.

### Supplementary Tables

**Table S1.** Posterior results from running DenMark on a subset of MERFISH genes.

| Parameter | *Aqp4* Mean (95% CI) | *Cxcl12* Mean (95% CI) | *Cxcr4* Mean (95% CI) | *Egfr* Mean (95% CI) | *Gfap* Mean (95% CI) |
|---|---|---|---|---|---|
| **Correlation** | | | | | |
| Latent field correlation ($\rho$) | 0.88 (0.80, 0.94) | 0.67 (0.52, 0.80) | 0.85 (0.74, 0.92) | 0.52 (0.16, 0.78) | 0.52 (0.43, 0.60) |
| **Intercepts** | | | | | |
| Average cells per unit area (exp($\beta_0$)) | 0.26 (0.01, 4.10) | 0.20 (0.01, 3.06) | 1.06 (0.08, 8.85) | 0.15 (0.01, 2.08 ) | 0.78 (0.08, 5.99) |
| Average expression per cell ($\exp(\beta_1)$) | 0.31 (0.13, 0.67) | 0.09 (0.03, 0.24) | 0.00 (0.00, 0.00) | 0.28 (0.16, 0.44) | 0.01 (0.00, 0.04) |
| **Spatial Parameter** | | | | | |
| Cell clustering process length of scale, mm ($\phi_1$) | 1.74 (1.41, 2.08) | 1.76 (1.43, 2.13) | 1.57 (1.27, 1.93) | 1.85 (1.51, 2.23) | 1.58 (1.32, 1.86) |
| Gene-specific process length of scale, mm ($\phi_2$) | 0.18 (0.13, 0.23) | 0.14 (0.11, 0.19) | 0.23 (0.17, 0.31) | 0.20 (0.15, 0.27) | 0.16 (0.13, 0.19) |
| Cell clustering process, SD ($\sigma_1$) | 4.28 (3.38, 5.28) | 4.30 (3.35, 5.43) | 3.59 (2.80, 4.58) | 4.32 (3.42, 5.43) | 3.74 (3.36, 5.45) |
| Gene-specific process, SD ($\sigma_2$) | 1.32 (1.01, 1.69) | 1.94 (1.53, 2.44) | 3.35 (2.39, 4.44) | 0.62 (0.47, 0.86) | 6.60 (5.79, 7.56) |
| **Computation Time (mins)** | 173.5 | 195.0 | 195.4 | 187.5 | 180.7 |
| **max Rhat** | 1.03 | 1.01 | 1.01 | 1.02 | 1.02 |

**Table S2.** Posterior summary on running DenMark on selected genes in 10x Xenium breast cancer.

| | *CDH1* | | | *POSTN* | | | *TRAC* | | |
|---|---|---|---|---|---|---|---|---|---|
| **Parameter** | DCIS | Invasive | Immune | DCIS | Invasive | Immune | DCIS | Invasive | Immune |
| **Correlation** | | | | | | | | | |
| Latent field correlation | 0.83 (0.76, 0.88) | 0.73 (0.66, 0.80) | -0.50 (-0.66, -0.30) | -0.04 (-0.24, 0.16) | 0.28 (0.13, 0.42) | -0.19 (-0.34, -0.03) | -0.15 (-0.46, 0.22) | -0.16 (-0.50, 0.25) | 0.40 (0.25, 0.53) |
| **Intercepts** | | | | | | | | | |
| Average cells per unit area ($mm^2$) | 0.03 (0.00, 0.28) | 0.68 (0.10, 3.71) | 41.26 (27.39, 61.56) | 0.06 (0.01, 0.46) | 0.80 (0.13, 4.18) | 40.85 (26.58, 61.56) | 0.05 (0.01, 0.45) | 1.01 (0.19, 4.76) | 38.86 (25.53, 58.56) |
| Average expression per cell | 0.85 (0.55, 1.26) | 0.42 (0.22, 0.72) | 0.21 (0.16, 0.28) | 0.23 (0.15, 0.33) | 0.17 (0.11, 0.27) | 1.31 (1.10, 1.55) | 0.09 (0.06, 0.12) | 0.08 (0.05, 0.12) | 2.77 (2.51, 3.06) |
| **Spatial Parameter** | | | | | | | | | |
| Length of scale (clustering process), mm | 0.25 (0.22, 0.28) | 0.33 (0.28, 0.37) | 0.24 (0.20, 0.29) | 0.24 (0.21, 0.27) | 0.33 (0.28, 0.38) | 0.24 (0.20, 0.29) | 0.24 (0.22, 0.27) | 0.32 (0.28, 0.38) | 0.25 (0.21, 0.30) |
| Length of scale (gene-specific process), mm | 0.24 (0.18, 0.32) | 0.21 (0.16, 0.27) | 0.38 (0.26, 0.55) | 0.20 (0.14, 0.26) | 0.16 (0.12, 0.22) | 0.23 (0.18, 0.29) | 0.20 (0.14, 0.27) | 0.36 (0.24, 0.50) | 0.26 (0.19, 0.35) |
| Standard Deviation (clustering process) | 13.3 (12.4, 14.4) | 7.45 (6.68, 8.27) | 2.37 (2.15, 2.63) | 13.0 (12.0, 14.1) | 7.17 (6.40, 8.00) | 2.40 (2.16, 2.66) | 13.0 (12.0, 14.0) | 7.09 (6.38, 7.90) | 2.43 (2.19, 2.70) |
| Standard Deviation (gene-specific process) | 2.1 (1.9, 2.4) | 2.66 (2.32, 3.03) | 0.92 (0.79, 1.07) | 2.0 (1.7, 2.3) | 2.33 (1.88, 2.92) | 0.94 (0.84, 1.06) | 1.08 (0.90, 1.29) | 0.86 (0.69, 1.05) | 0.45 (0.39, 0.52) |
| **Computation Time (mins)** | 151.3 | 136.59 | 66.05 | 271.8 | 131.9 | 76.3 | 148.7 | 133.0 | 67.7 |
| **max Rhat** | 1.02 | 1.03 | 1.00 | 1.02 | 1.02 | 1.00 | 1.02 | 1.01 | 1.00 |

#### The list of density-correlated genes identified from Mouse Brain MERFISH Data

See Data S1.

#### The list of density-correlated genes identified from Breast Cancer 10X Xenium data

See Data S2.
